# Machine learning approach to identify resting-state functional connectivity pattern serving as an endophenotype of autism spectrum disorder

**DOI:** 10.1101/348599

**Authors:** Bun Yamagata, Takashi Itahashi, Junya Fujino, Haruhisa Ohta, Motoaki Nakamura, Nobumasa Kato, Masaru Mimura, Ryu-ichiro Hashimoto, Yuta Aoki

**Author notes:** These authors equally contributed to this work (Bun Yamagata, Takashi Itahashi). **Corresponding author:** Yuta Aoki, MD, PhD, Senior Assistant Professor, Medical Institute of Developmental Disabilities Research at Showa University, 6-11-11 Kitakarasuyama, 157-8577, Tokyo, Japan.

## Abstract

Endophenotype refers to a measurable and heritable component between genetics and diagnosis and exists in both individuals with a diagnosis and their unaffected siblings. We aimed to identify a pattern of endophenotype consisted of multiple connections. We enrolled
adult male individuals with autism spectrum disorder (ASD) endophenotype (i.e., individuals with ASD and their unaffected siblings) and individuals without ASD endophenotype (i.e., pairs of typical development (TD) siblings) and utilized a machine learning approach to
classify people with and without endophenotypes, based on resting-state functional connections (FCs). A sparse logistic regression successfully classified people as to the endophenotype (area under the curve=0.78, classification accuracy=75%), suggesting the existence of endophenotype pattern. A binomial test identified that nine FCs were consistently selected as inputs for the classifier. The least absolute shrinkage and selection operator with these nine FCs predicted severity of communication impairment among individuals with ASD (*r*=0.68, *p*=0.021). In addition, two of the nine FCs were statistically significantly correlated with the severity of communication impairment (*r*=0.81, *p*=0.0026 and *r*=-0.60, *p*=0.049). The current findings suggest that an ASD endophenotype pattern exists in FCs with a multivariate manner and is associated with clinical ASD phenotype.

## Introduction

Autism spectrum disorder (ASD) is a developmental disorder characterised by deficits in social interaction and repetitive restricted behavior (American Psychiatric Association, 2013). Examples of these social interaction impairments among people with ASD include theory of mind, empathy, and facial emotion recognition (Simon Baron-Cohen, Leslie, & Frith, 1985; S. Baron-Cohen & Wheelwright, 2004; Uljarevic & Hamilton, 2013). Consistent with the findings that such social interactions depend on the social brain neural system (Adolphs, 2009), abnormalities within the social brain have been observed among individuals with ASD (Yuta Aoki, Cortese, & Tansella, 2015; Kana, Keller, Cherkassky, Minshew, & Just, 2009; Kleinhans, Richards, Greenson, Dawson, & Aylward, 2016; Murphy et al., 2017; Pelphrey, Shultz, Hudac, & Vander Wyk, 2011). Besides the anatomical extension of abnormalities, individuals with ASD consistently present atypical functional connections (FCs) (Cherkassky, Kana, Keller, & Just, 2006; Di Martino et al., 2014; Uddin, Supekar, & Menon, 2013). Thus, atypical FCs within the social brain are of interest and may underlie the impairment in social interactions among individuals with ASD.

ASD is a highly heritable condition (Colvert et al., 2015). Indeed, although the prevalence of ASD is about 1% in the general population (Autism & Developmental Disabilities Monitoring Network Surveillance Year Principal, 2014), the risk of developing ASD increases by up to 20% among biological siblings of individuals with ASD (Ozonoff et al., 2011). Despite such obvious heritability, the concordance of clinical diagnoses among monozygotic twins is only about 60% indicates, indicating that ASD is influenced by complex genetic interactions (Hallmayer et al., 2011). Reflecting this complex genetic contribution, autistic traits have a continuous distribution among people with incomplete genetic traits of ASD (Constantino & Todd, 2003). In fact, biological family members of individuals with ASD often have subclinical difficulties in social interactions (Piven & Palmer, 1999).

Endophenotype refers to a measurable and heritable component between genes and disease diagnosis (Gottesman & Gould, 2003). Because of the complex genetic contribution, characterising the ASD endophenotype is particularly important because it may provide an objective intermediate marker and insight into the pathophysiology of ASD. Reflecting its heritability and consistently observed abnormalities among individuals with ASD (Di Martino et al., 2014; Glahn et al., 2010), resting-state functional magnetic resonance imaging (R-fMRI) is one of promising modalities that quantify ASD endophenotype (Khadka et al., 2013).

To identify the ASD endophenotype, prior neuroimaging studies using several modalities have enrolled individuals with ASD, their unaffected siblings, and typical development (TD) (Barnea-Goraly, Lotspeich, & Reiss, 2010; Jou et al., 2016; Moseley et al., 2015). These studies separately compared each connection across these three groups and showed the shared atypical findings among unaffected siblings of individuals with ASD and individuals with ASD. Most of prior studies recognised such shared abnormality, which is widely scattered throughout the brain, as an endophenotype.

The aim of this study was three-folds. First, we aimed to classify pairs of people as to the ASD endophenotype using a multivariate machine learning approach. Motivation of applying a machine learning approach is that we recognised that ASD endophenotype likely consisted of multiple FCs with different weights (i.e., as a pattern of altered FCs), rather than an unweighted integration of abnormalities shared by individuals with ASD and their unaffected siblings. Second, we examined which FCs were consistently selected when classifying people as to the endophenotype. Finally, to confirm the relation between the selected FCs and the clinical ASD phenotype, we employed the least absolute shrinkage and selection operator (LASSO), to predict the severity of social interaction impairment among people with ASD using the FCs serving as the endophenotype. To do so, we obtained R-fMRI data from 60 participants, consisting of 30 people with ASD endophenotype (15 individuals with ASD and 15 of their unaffected siblings) and 30 people without the ASD endophenotype (15 pairs of TDs). Since ASD is characterized by the impairment of social interactions, we focused on FCs within the social brain. To increase the homogeneity of the participants, we enrolled age- and IQ-matched males.

## Materials and methods

### Participants

We analysed data from 60 adult males consisting of 30 pairs of biological siblings. Thirty people had the ASD endophenotype. Specifically, 15 pairs of participants were discordant for the diagnosis of ASD: namely, one of the siblings was affected with ASD, and the other was unaffected. Another 30 people did not have the endophenotype and consisted of 15 pairs of TD siblings. None of the TD siblings had a family member who had been diagnosed as having ASD. All the participants with ASD were diagnosed by experienced psychiatrists based on the DSM-IV-TR (American Psychiatric, 2000). The diagnosis was further supported by the Autism Diagnostic Observation Schedule (ADOS) (Lord et al., 2001). In addition, to confirm the absence of a diagnosis of ASD in the unaffected sibling, the parents of the siblings discordant for the diagnosis of ASD were interviewed using the Autism Diagnostic Interview-Revised (ADI-R) (Tsuchiya et al., 2013). We used the Edinburgh Handedness Inventory to evaluate handedness (Oldfield, 1971). The intelligence quotient (IQ) of each participant was assessed using either the Wechsler Adult Intelligence Scale-Third Edition or the WAIS-Revised (Wechsler, 1997; Wechsler & De Lemos, 1981). All the participants completed the Japanese version of the Autism-Spectrum Quotient (AQ) (Wakabayashi, Baron-Cohen, Wheelwright, & Tojo, 2006). Psychiatric comorbidities were observed in two participants with ASD: one had attention-deficit/hyperactivity disorder, and the other had learning disabilities. Five of the participants were taking medication at the time of scanning: benzodiazepine (*n* = 3), anti-depressants (*n* = 3), and psychostimulant (*n* = 1). For inclusion in the TD group, we confirmed the absence of an ASD diagnosis among the family member and the absence of an Axis I diagnosis per the DSM-IV-TR using the Mini-International Neuropsychiatric Interview (Hergueta, Baker, & Dunbar, 1998). The absence of a history of psychotropic medication use was also required. The exclusion criteria for all the participants were known genetic diseases, an estimated full intelligence quotient (IQ) of 80 or below, or the total AQ score of 33 or above. Written informed consent was obtained from all the participants, after they had received a complete explanation of the study. The Ethics Committee of Showa University approved the study protocol. The study was prepared in accordance with the ethical standards of the Declaration of Helsinki.

### MRI acquisition

All MRI data were acquired using a 3.0 T MRI scanner (MAGNETON Verio, Siemens Medical Systems, Erlangen, Germany) with a 12ch head coil. Functional images were acquired with an echo planar imaging sequence (repetition time [TR]: 2500 ms, echo time [TE]: 30 ms, flip angle: 80°, field of view [FOV]: 212 mm, matrix size: 64 × 64, slice thickness: 3.2 mm with a 0. 8-mm gap, 40 axial slices) at rest for 10 min 10 s (244 volumes). During the resting-state scans, participants were asked to gaze at a cross-hair displayed at the center of the screen, not to think about specific things, and to stay awake. To correct for the distortion of the functional image, gradient echo field mapping images were acquired immediately after the resting-state scans (TR: 488 ms, short TE: 4.92 ms, long TE: 7.38 ms, flip angle: 60°, FOV: 212 mm, matrix size: 64 × 64, slice thickness: 3.2 mm with a 0.8-mm gap, 40 axial slices). For normalisation purpose, a T1-weighted image was acquired using an MPRAGE sequence (TR: 2.3 s, TE: 2.98 ms, flip angle: 9°, FOV: 256 mm, matrix size: 256 × 256, slice thickness: 1mm, 240 sagittal slices, voxel size: 1×1×1 mm).

### R-fMRI data preprocessing

All the functional images were preprocessed using the Statistical Parametric Mapping (SPM12; Wellcome Department of Cognitive Neurology, London, UK) and functions implemented in FMRIB’s Software Library (FSL; https://fsl.fmrib.ox.ac.uk/fsl/fslwiki/). Preprocessing was performed as follows: 1) the first four volumes were discarded to allow for T1 equilibration; 2) slice timing correction; 3) head motion correction using mcflirt implemented in the FSL (Jenkinson, Bannister, Brady, & Smith, 2002), 4) distortion correction using the FUGUE implemented in the FSL, 5) co-registration of functional images to an anatomical image, 6) spatial normalisation and resampling to a resolution of 2 × 2 × 2 mm, and 7) spatial smoothing (6-mm full-width at half-maximum).

To remove the effects of subtle head motions during the scans (Power, Barnes, Snyder, Schlaggar, & Petersen, 2012), ICA-AROMA was applied (Pruim, Mennes, Buitelaar, & Beckmann, 2015; Pruim, Mennes, van Rooij, et al., 2015). After ICA-AROMA was performed, nuisance regression was further performed. Nuisance signals consisted of signals averaged over white matter, cerebrospinal fluid, and grey matter, respectively (Parkes, Fulcher, Yu Cel, & Fornitod, 2017; Yahata et al., 2016). A band-pass filter (0.008 - 0.1 Hz) was then applied to the residual time-series in a voxel-wise manner. For each participant, the mean frame-wise displacement (FD) was calculated from head motion parameters to quantify the amount of head motion during scans (Jenkinson et al., 2002). The mean FD of each group is shown in Table 1.

**Table 1:**
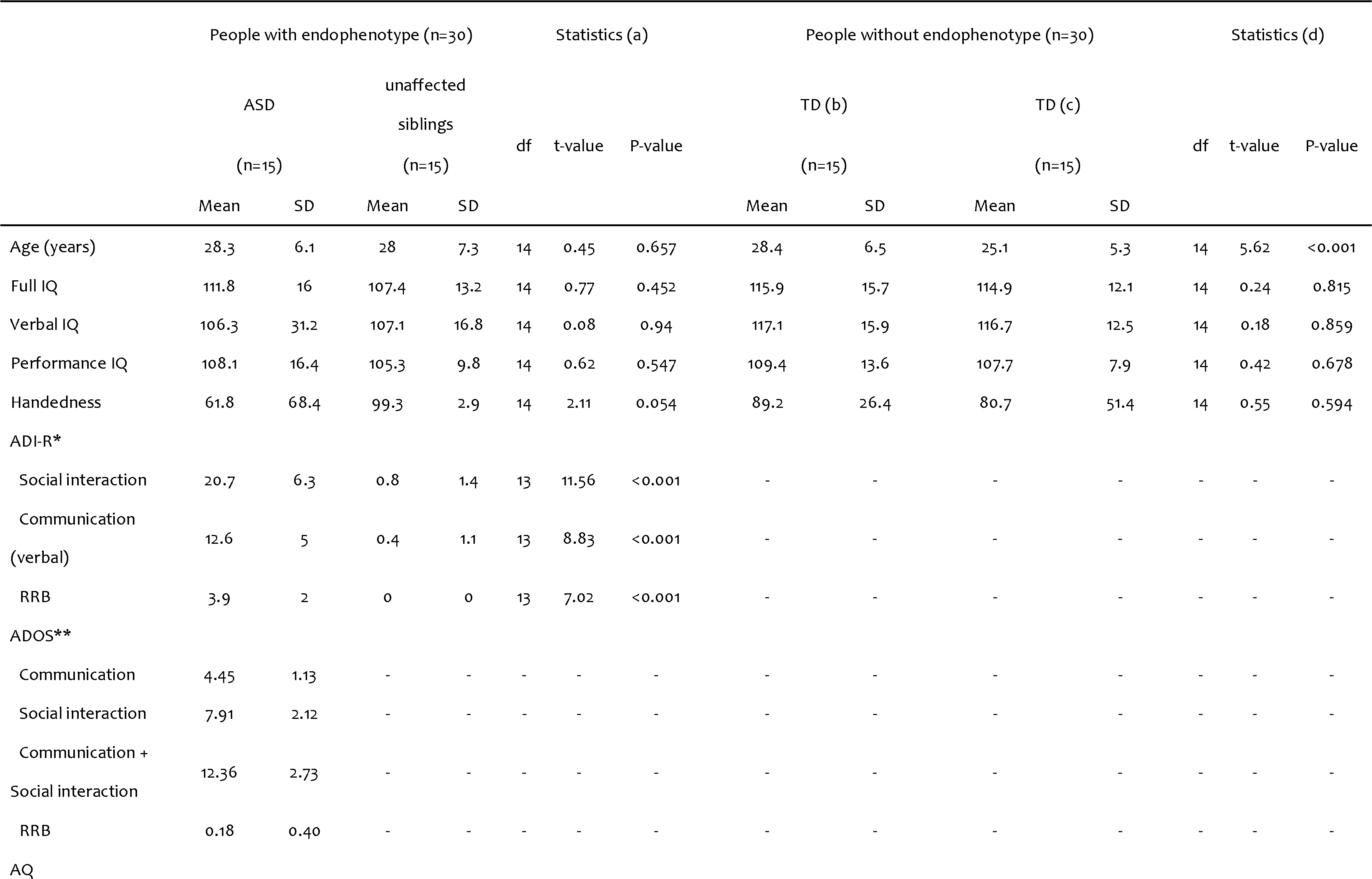
Characteristics of the participants.

|  |  |  |  |  |  |  |  |  |  |  |  |  |  |  |
| --- | --- | --- | --- | --- | --- | --- | --- | --- | --- | --- | --- | --- | --- | --- |
| Total | 34.2 | 6.2 | 19.4 | 6.9 | 12 | 6.27 | <0.001 | 17.6 | 5.5 | 14.9 | 6 | 14 | 1.12 | 0.28 |
| AS | 7.4 | 1.6 | 3.9 | 1.8 | 12 | 6.3 | <0.001 | 4 | 2.1 | 3.7 | 1.4 | 14 | 0.54 | 0.596 |
| ATD | 5.4 | 1.9 | 3.2 | 1.9 | 12 | 2.67 | 0.02 | 3.4 | 2.1 | 3.9 | 1.6 | 14 | 0.88 | 0.396 |
| COM | 7.6 | 1.6 | 3.6 | 2.6 | 12 | 4.16 | 0.001 | 3.1 | 2.5 | 2.6 | 2.3 | 14 | 0.5 | 0.625 |
| IMG | 6 | 2 | 4.1 | 1.4 | 12 | 2.31 | 0.04 | 3.7 | 1.7 | 2.8 | 1.7 | 14 | 1.26 | 0.229 |
| SS | 7.9 | 2 | 4.5 | 2.7 | 12 | 4.26 | 0.001 | 3.4 | 2.7 | 2 | 2.1 | 14 | 1.38 | 0.191 |
| SES (e) | 5.5 | 1.1 | 5.5 | 1.1 | 14 | 0.2 | 0.843 | 5.9 | 1.2 | 5.7 | 1.1 | 13 | 0.46 | 0.655 |
| Mean FD | 0.17 | 0.07 | 0.14 | 0.05 | 14 | 1.51 | 0.153 | 0.16 | 0.07 | 0.18 | 0.10 | 14 | 0.687 | 0.503 |

(a) Statistics show the results of comparisons between adult males with ASD and their unaffected brothers. (b) Older siblings. (c) Younger siblings. (d) Statistics show the results of comparisons between typical older and younger siblings. (e) A higher score indicates a lower socioeconomic status (Okada et al., 2014). Abbreviations: ADOS-2: Autism Diagnostic Observation Schedule Second Edition, ADI-R: Autism Diagnostic Interview-Revised, AQ: autism-spectrum quotient, AS: attention switching/tolerance of change, ASD: autism spectrum disorder, ATD: attention to detail, COM: communication skills, IMG: imagination, IQ: intelligence quotient, RRB: restricted repetitive behaviors, SD: standard deviation, SES: socioeconomic status, SS: social skills, TD: typical development. \*The ADI-R score was missing for one person. \*\*ADOS scores are missing for four people.

The social brain connectome atlas was used for network construction (Alcala-Lopez et al., 2017). This atlas consists of 36 brain regions associated with social functioning, such as theory of mind, empathy, and facial emotion recognition. For each participant, the mean time-series was extracted from each region of interest (ROI) and the correlation coefficients between all possible pairs of ROIs were calculated, resulting in a 36 × 36 correlation matrix for each participant. Finally, Fisher’s *r*-to-*z* transformation was applied to each correlation coefficient This procedure generated 630 features for each participant.

### Statistical analysis

#### Identification of endophenotype pattern of FCs

Since our aim was to identify a pattern of FCs serving as an endophenotype from a given data, we formulated the problem in the following manner. Based on previous findings that individuals with ASD and their unaffected siblings shared a pattern of alterations when compared to TDs (Moseley et al., 2015), we recognised that endophenotype was a measurable component satisfying the following conditions:

1. individuals with ASD have higher values than TDs (i.e., ASD > TD);
2. unaffected siblings of individuals with ASD also have higher values than TDs (i.e., unaffected sibling > TD);
3. TD sibling pairs have similar values (i.e., TD ? TD sibling).

According to the conditions (1) and (2), our aim could be achieved by solving a classification problem, in which we intended to identify FCs that could discriminate persons with a high endophenotype value from persons with a low endophenotype value. Individuals with ASD and their unaffected siblings were regarded as having the ASD endophenotype, while TD and TD siblings were regarded as not having the endophenotype throughout this manuscript.

To examine whether the FCs serve as an endophenotype, we employed sparse logistic regression (SLR) (Yamashita, Sato, Yoshioka, Tong, & Kamitani, 2008). SLR can train a logistic regression model while automatically selecting endophenotype-related FCs. Briefly, SLR relies on a hierarchical Bayesian estimation, in which the prior distribution of each element of the parameter vector is represented as a Gaussian distribution. Based on the automatic relevance determination, irrelevant features are not used in the classification because the respective Gaussian prior distributions have a sharp peak at zero. Such efficient feature elimination method implemented in SLR can mitigate the problems of over-fitting caused by a small sample size. To evaluate the performance of the classifier, a leave-one-pair-out cross-validation (LOPOCV) was performed. Of note, the term ‘pair’ stands for sibling pairs. In each fold, all-but-one pair was used to train the SLR classifier, while the remaining pair was used for the evaluation.

To further examine the statistical significance of the classification accuracy, a permutation test was performed. At each iteration, a permuted dataset was generated by shuffling the endophenotype label while keeping the pair information. Then, LOPOCV was performed to calculate the classification accuracy for the permuted dataset. This procedure was repeated 5,000 times to construct a null distribution. Statistical significance was set at *P* < 0.05.

#### Binomial test

SLR selects a small number of relevant FCs from a given data. To confirm that FCs selected by the classifier were not randomly selected, the statistical significance of the selection counts was examined using a binomial test. The classifier selected 8.13 ± 0.94 (mean ± standard deviation [SD]) out of 630 FCs in each of the 30 validation folds (see Results). Thus, we assumed a binomial distribution, *Bi*(*n*, *p*), where *n* stands for the number of validation folds (i.e., *n* = 30) and *p* stands for the probability of being selected from the set of FCs (i.e., *p* = 8/630).

#### Relation between endophenotype-related FCs and clinical phenotype

Once the endophenotype-related FCs had been identified by the classifier with binomial tests, we predicted the severity of the clinical symptoms measured by the ADOS. The score of each individual was predicted using LASSO with nine FCs consistently selected in the classifier (Tibshirani, 1996). While an optimal regularization parameter in the LASSO was determined by an internal 10-fold CV, the weight parameters were determined through a leave-one-subject-out CV (LOSOCV), in which the scores of all-but-one participants were linearly regressed using the nine FCs as explanatory variables. Given the notion that age-related differences might have an impact on the severity of clinical symptoms (Lee, Park, James, Kim, & Park, 2017), we added age as an additional explanatory variable. To evaluate how predicted and actual scores matched, the correlation coefficient was calculated. We further examined the extent to which the FCs were significantly correlated with the severity of clinical symptoms by calculating correlation coefficient. Because of the explorative of this analysis nature, we used a liberal statistical threshold of *P* < 0.05.

## Results

### Identification of endophenotype pattern of FCs

We discriminated participants with the ASD endophenotype (i.e., individuals with ASD and their unaffected siblings) from TDs using an SLR with the LOPOCV. This classifier separated participants with endophenotype from TDs with 75% accuracy (sensitivity = 76.67% and specificity = 73.33%) and an area under the curve (AUC) of 0.78 (permutation test with 5,000 iterations, *p* < 0.001; Figures la and lb), suggesting that FCs selected by this classifier captured the endophenotype-related features.

**Figure 1.**
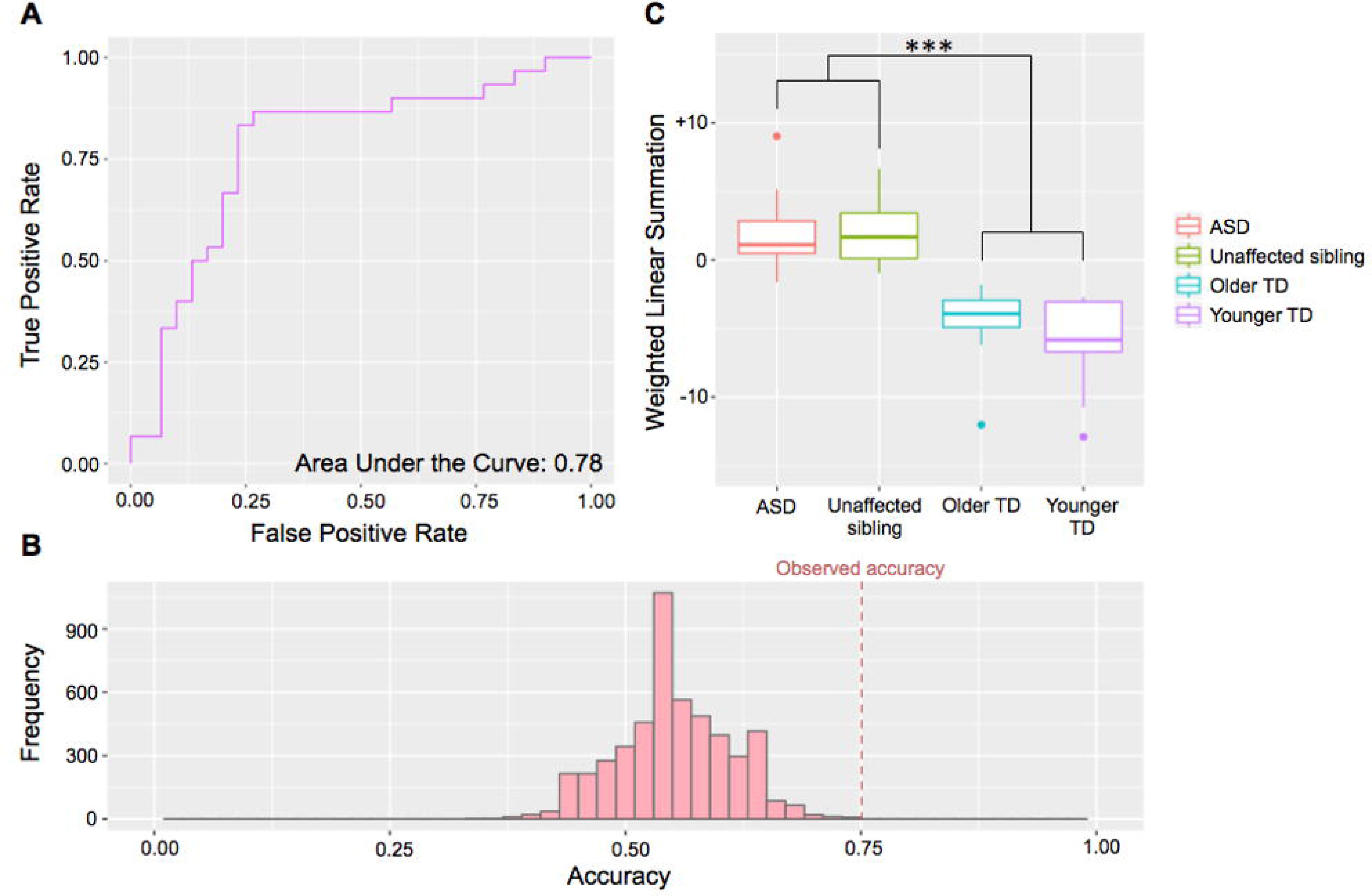
Classification results and post-hoc tests. (a) Receiver operating characteristic curve. A sparse logistic regression with leave-one-pair-out cross validation exhibited an area under the curve of 0.78. (b) Null distribution of classification accuracy derived by a permutation test with 5,000 iterations. The permutation test demonstrated that the observed classification accuracy (75%) was significantly higher than that obtained by the permuted classifier (*p* < 0.001). (c) Post-hoc paired t-tests demonstrated that there were no statistically significant differences between individuals with autism spectrum disorder (ASD) and their unaffected siblings in the weighted linear summation of selected FCs (*t*-value = −0.03, *df* = 14, *p* = 0.98) as well as between TD pairs (*t*-value = 1.14, *df* = 14, *p* = 0.27). On the other hand, persons with endophenotype exhibited significantly higher values than those without endophenotype (*t*-value = 10.28, *df* = 58, *p* < 0.001). ***: *P* < 0.001.

Post-hoc paired *t*-tests were performed to examine whether the conditions (see Methods) were satisfied. As shown in Figure 1c, paired *t*-tests demonstrated that the weighted linear summation (WLS) of FCs selected in the classifier was not statistically significantly different between individuals with ASD and their unaffected siblings (*t*-value = −0.03, *df* = 14, *p* = 0.98). In contrast, a two-sample *t*-test showed statistically significant differences between people with endophenotype (i.e., individuals with ASD and their unaffected siblings) and those without endophenotype (*t*-value = 10.28, *df* = 58, *p* < 0.001). These results indicate that the WLS of FCs selected by the classifier satisfied the set of conditions regarding endophenotype, but not the clinical diagnosis. Furthermore, the analysis did not show any significance difference between TD sibling pairs (*t*-value = 1.14, *df* = 14, *p* = 0.27).

To rule out the possibility that nuisance covariates (i.e., head motion and age) have potential to be important features for classification, we repeated the same classification analysis while adding the two nuisance variables as additional explanatory variables (i.e., 630 FCs + 2 nuisance variables). If the nuisance variables could explain the endophenotype label over the previously selected FCs, then the SLR would automatically select these nuisance variables as input for the logistic function. We confirmed that the classification accuracy and the AUC were not changed by adding these nuisance variables (accuracy = 75% and AUC = 0.78). Furthermore, in each fold, the SLR never selected the two nuisance variables as inputs for the logistic function (i.e., the weighted parameters for these variables were zero).

### Binomial test

Furthermore, we investigated which FCs were stably selected by the SLR across the LOPOCV. In each fold, the classifier selected 8.13 ± 0.94 (mean ± SD) out of the 630 FCs across 30 validation folds. We counted how many times each FC was selected. Under the null hypothesis that eight FCs were randomly selected from 630 FCs, a binomial test was applied to examine the probability of the selection count. We found that nine FCs were selected at a significant frequency (*p* < 0.05, Bonferroni corrected for 630 connections; Figure 2 and Table 2). The selection count for these nine FCs was 22.78 ± 7.61 while that for the remaining FCs was 0.0627 ± 0.33. This result indicates that the nine FCs were consistently selected across the validation folds.

**Figure 2.**
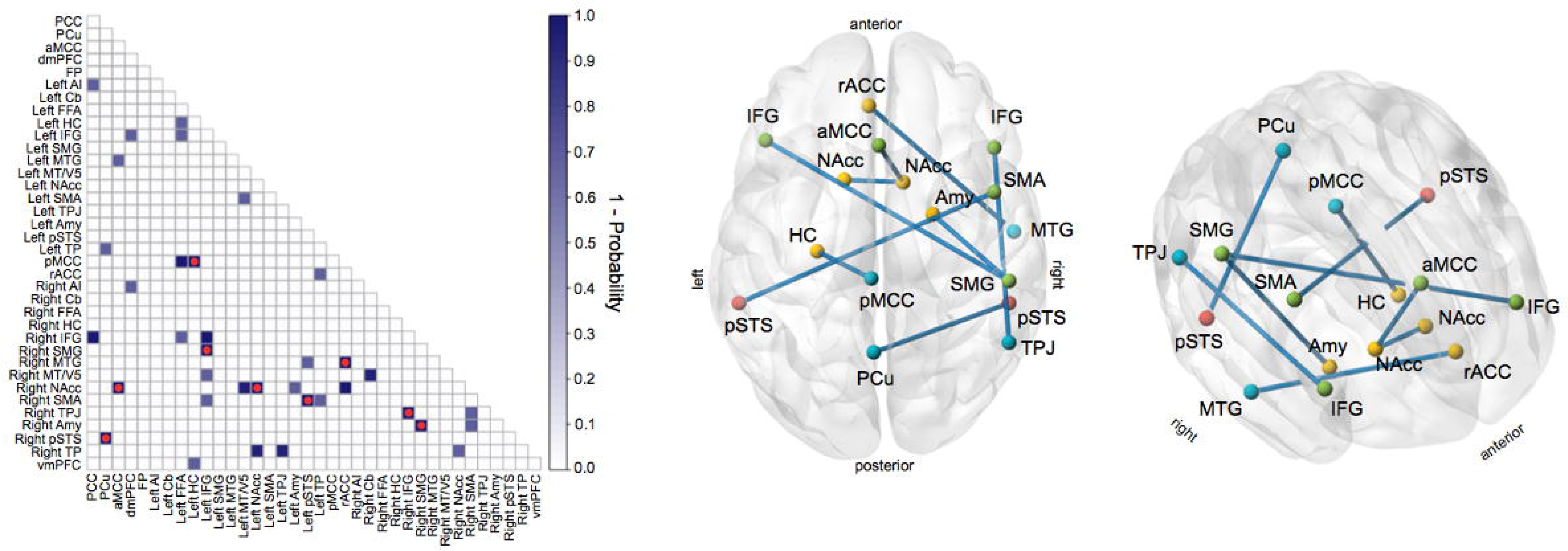
Results of identified functional connections (FCs) stably selected for the classification. The left panel shows the results of binomial tests. The red circle indicates the statistical significance of the selected count after multiple comparison correction. The right panel displays the distribution of nine FCs exhibiting statistical significance after multiple comparison corrections. The colors of nodes represent the sub-systems within the social brain atlas defined in (Alcala-Lopez et al.) Red: visual-sensory system, Yellow: limbic system, Green: intermediate-level system, Cyan: higher-level system. **Abbreviations:** AI: anterior insula, aMCC: anterior middle cingulate cortex, Amy: amygdala, Cb: cerebellum, dmPFC: dorsomedial prefrontal cortex, FFA: fusiform face area, FP: frontal pole, HC: hippocampal cortex, IFG: inferior frontal gyrus, MT/V5: middle temporal V5 area, MTG: middle temporal gyrus, NAcc: nucleus accumbens, PCC: posterior cingulate cortex, PCu: precuneus, pSTS: posterior superior temporal sulcus, SMA: supplementary motor area, SMG: supramarginal gyrus, TPJ: temporo-parietal junction, vmPFC: ventromedial prefrontal cortex

**Table 2:**
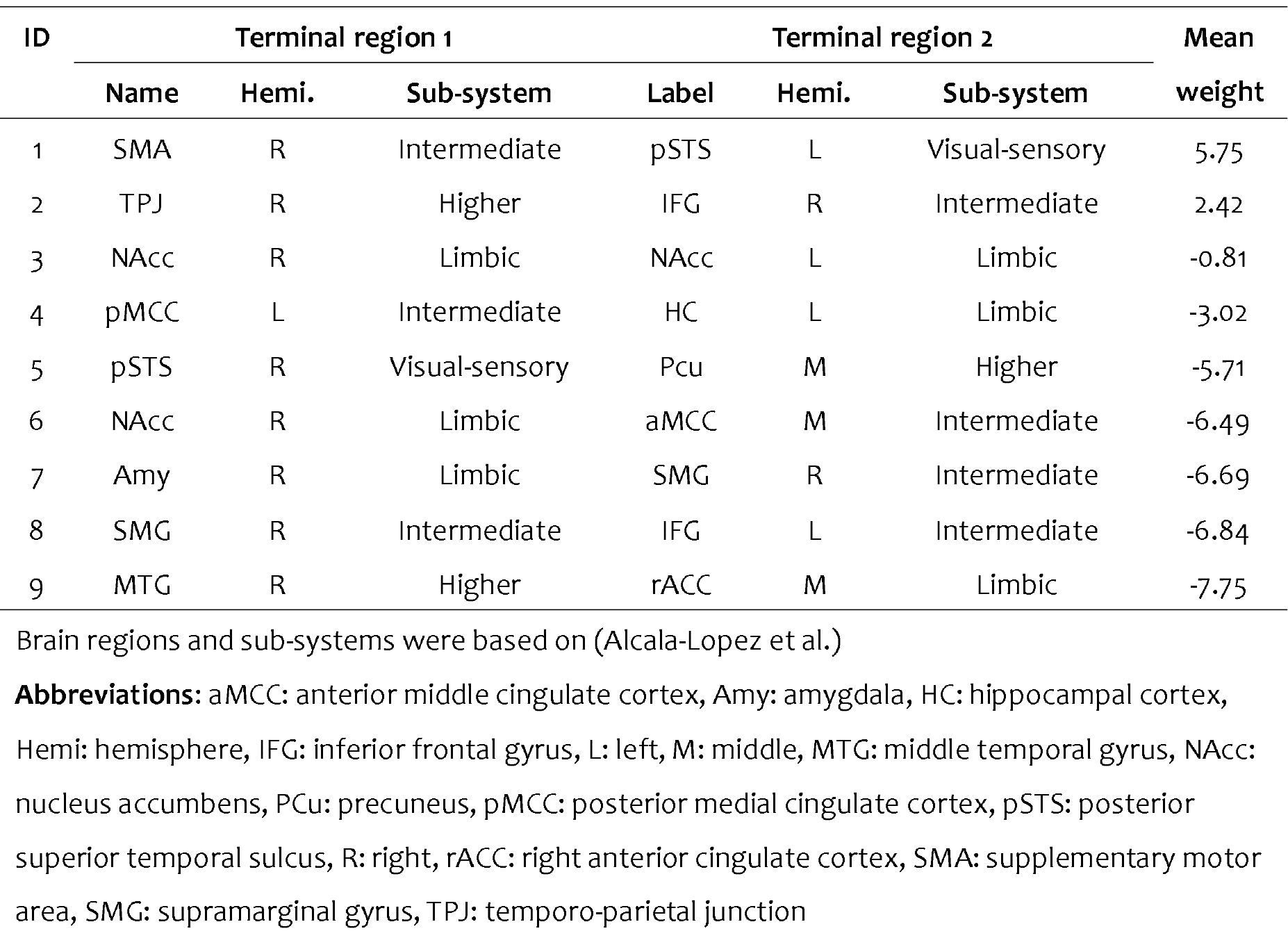
The list of functional connections (FCs) stably selected for classification.

### Relation between endophenotype-related FCs and clinical phenotype

We predicted the severity of the clinical symptoms measured by the ADOS using the nine FCs identified in the classifier for endophenotype. LASSO with LOSOCV was individually applied to determine the weights of the nine FCs so that their weighted linear summation was used as a predictor for the corresponding severity score. We found that the communication domain of the ADOS was well predicted from the nine FCs with a statistically significant correlation (*r* = 0.68, *p* = 0.021; Figure 3a). Furthermore, we found that two of nine FCs were significantly correlated with the severity of impaired communication measured by ADOS: one was FC between the right temporo-parietal junction and right inferior frontal gyrus (*r* = 0.81, *p* = 0.0026; Figure 3b), and the other was FC between the right nucleus accumbens and anterior middle cingulate cortex (*r* = −0.60, *p* = 0.049; Figure 3c).

**Figure 3.**
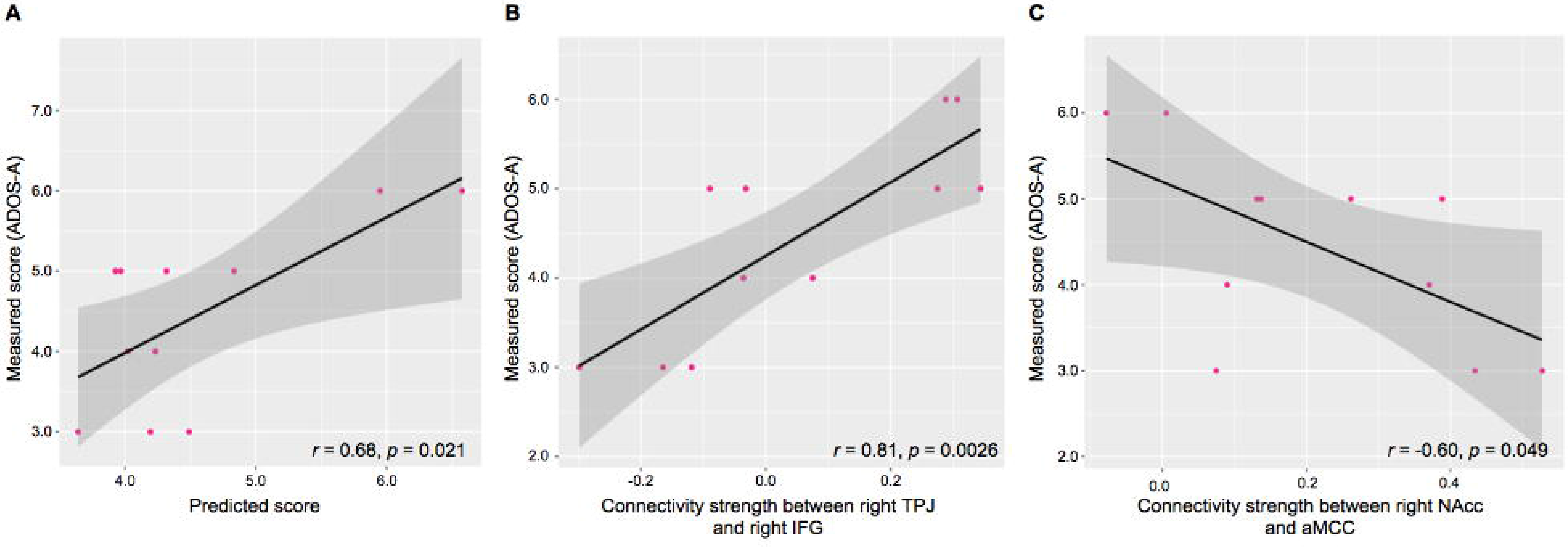
Associations between the severity of clinical symptoms and functional connectivity (FCs) selected by the classifier. (a) Prediction of the severity of communication measured by the Autism Diagnostic Observation Schedule (ADOS) using the least absolute shrinkage and selection operator with leave-one-subject-out cross-validation. The correlation coefficient between the predicted and measured scores exhibited a significant positive correlation (*r* = 0.68, *p* = 0.021). (b) The connectivity strength between the right temporo-parietal junction and right inferior frontal gyrus was significantly correlated with the severity of impaired communication as measured by the ADOS-A (*r* = 0.81, *p* = 0.0026). (c) The connectivity strength between the right nucleus accumbens and anterior middle cingulate cortex was significantly correlated with the severity of communication impairment as measured by the ADOS-A (*r* = −0.60, *p* = 0.049).

## Discussion

Our novel framework was motivated by 1) the observation that neural correlates of ASD emerge as a pattern of FCs and 2) the lack of any previous study attempting to identify the endophenotype as a pattern of FCs. Thus, we applied the machine learning approach to classify people as to the ASD endophenotype. Subsequent binomial tests identified nine FCs serving as the endophenotype. The LASSO further revealed that the selected FCs were associated with the clinical phenotype among individuals with ASD. This high-performance proof-of-concept study shows the importance and relevance of using the machine learning approach to identify the endophenotype as a pattern of FCs.

We utilised the SLR to classify participants as to the ASD endophenotype. Most prior neuroimaging studies identified abnormalities shared by individuals with ASD and their unaffected siblings or abnormalities in the siblings that were intermediate between ASD and TD as an endophenotype (Barnea-Goraly et al., 2010; Jou et al., 2016; Moseley et al., 2015). In these studies, each variable (i.e., connectivity index) was separately examined to determine whether it belonged to the endophenotype. In contrast, we applied a multivariate and data-driven approach to classify individuals with or without the ASD endophenotype using all the available variables (i.e., 630 FCs within the social brain). This approach inherently assumes that the endophenotype emerges as a pattern of altered FCs and allows all the variables to contribute to the endophenotype to different extents. In the case of ASD, the genetic abnormalities vary across individuals (Geschwind & Levitt, 2007; Miles, 2011). In addition, multiple FCs are altered (Di Martino et al., 2014; Uddin et al., 2013). Thus, the genetic influence is not identical across FCs (Ameis & Szatmari, 2012; Meyer-Lindenberg & Weinberger, 2006; Zhan et al., 2014), and the endophenotype should emerge as multiple FCs with different extents. The consistency with the definition of the endophenotype might explain the good performance of classification.

The current study identified an ASD endophenotype among the FCs within the social brain. Since not all individuals with the endophenotype develop ASD clinical diagnosis (Ozonoff et al., 2011), the neural correlates for the endophenotype and the diagnosis may differ. Although we did not investigate the neural correlates for the diagnosis because of the small sample size, neural correlates for the clinical diagnosis of ASD might be located within the social brain (Yuta Aoki et al., 2015; Kana et al., 2009; Kleinhans et al., 2016; Murphy et al., 2017; Pelphrey et al., 2011). Given that there was no significant difference in the WLS of the selected FCs between individuals with ASD and their unaffected siblings, the selected FCs represent the ASD endophenotype, rather than the ASD diagnosis. Although it is beyond the scope of this proof-of-concept study, future research involving a large sample size is expected to dissociate the neural correlates of the ASD clinical diagnosis and the ASD endophenotype.

Using the LASSO, the selected FCs successfully predicted the severity of communication deficits among individuals with ASD. This relationship corroborates the assumption that the selected FCs are associated with the pathophysiology of ASD. Since we lacked any ADOS scores from the unaffected siblings of individuals with ASD, this relationship was observed only among the individuals with ASD. However, even if the ADOS scores were available from the unaffected siblings, the scores might not reflect the distribution of behaviour well, since the ADOS aims to measure clinical symptoms, not subclinical ones observed among people with the ASD endophenotype (Toth, Dawson, Meltzoff, Greenson, & Fein, 2007). Future studies enrolling unaffected siblings should obtain both MRI data and psychological evaluations that reflect the differences in ASD traits among subclinical individuals.

Several limitations should be considered in this study. First, although we successfully added the condition that the FCs should not differ between TD siblings (see Method), which was not possible in prior studies with only one TD group, the current study had a small sample size because of practical difficulties in recruitment. Thus, because of the lack of statistical power, we could not address brain regions outside of the social brain. However, brain regions outside the social brain are also likely to be involved in the endophenotype of ASD. In addition, although the sparseness of FC selection by the SLR mitigates over-fitting to the current sample, the current sample size was relatively small, compared with the number of FCs. A future study with a sample size large enough to avoid over-fitting is needed to examine the whole brain. Second, although the machine learning approach successfully classified individuals with or without the ASD endophenotype, the endophenotype may not be categorical, as with ASD diagnosis (Y. Aoki et al., 2017). Research using a large sample is encouraged to stratify individuals with an ASD endophenotype. Third, to increase the homogeneity of participants, we recruited only adult males in the current study. Given that the ASD brain shows an atypical developmental trajectory (Y. Aoki, Kasai, & Yamasue, 2012), the current finding may not be generalised to other age ranges. In addition, atypical sex differences have been observed among individuals with ASD (Lai et al., 2017). Thus, further studies that include only female ASD participants are expected.

## Conclusion

Using the machine learning approach, we demonstrated that the ASD endophenotype emerges as a pattern of FCs within the social brain. The FCs selected as inputs for the classifier were associated with the clinical ASD severity.

## Funding

This study is the result of “Development of BMI Technologies for Clinical Application” carried out under the Strategic Research Program for Brain Sciences by Japan Agency for Medical Research and Development (AMED). This work is partly supported by the grant by The Japan Foundation for Pediatric Research (to YA).

## Conflict of Interest

All authors report no conflict of interest.

## Ethical approval

The study was prepared in accordance with the ethical standards of the Declaration of Helsinki.

## Informed consent

Written informed consent was obtained from all the participants, after they had received a complete explanation of the study.

